# Speed and diffusion of kinesin-2 are competing limiting factors in flagellar length control model

**DOI:** 10.1101/751990

**Authors:** R M, NL H, WF M, H Q

## Abstract

Flagellar length control in *Chlamydomonas* is a tractable model system for studying the general question of organelle size regulation. We have previously proposed that diffusive return of the kinesin motor that powers intraflagellar transport can play a key role in length regulation. Here we explore how the motor speed and diffusion coefficient for the return of kinesin-2 affect flagellar growth kinetics. We find that the system can exist in two distinct regimes, one dominated by motor speed and one by diffusion coefficient. Depending on length, a flagellum can switch between these regimes. Our results indicate that mutations can affect length in distinct ways. We discuss our theory’s implication for flagellar growth influenced by beating and provide possible explanations for the experimental observation that a beating flagellum is usually longer than its immotile mutant. These results demonstrate how our simple model can suggest explanations for mutant phenotypes.

**Statement of Significance:** The eukaryotic flagellum is an ideal case study in organelle size control because of its simple linear shape and well-understood building mechanism. In our previous work, we proved that flagellar length in the green algae Chlamydomonas can be controlled by the diffusive gradient of the kinesin-2 motors that deliver building blocks to the tip. In this study, we expand on the analytical formulation of the diffusion model to show how physical parameters affect final length and regeneration time, enhancing the model’s potential to explain length mutants and motivate future research with precise predictions.

## 1. Introduction

Biologists have long been trying to understand how cells build themselves. The proteins that cells synthesize have to come together to form massive organized structures without any guidance. A striking case of this is that some single-celled organisms can regenerate missing pieces, implying that the cell has some form of design specifications embedded within it that allows the cell to reconstruct the correct form. The single-celled algae *Chlamydomonas reinhardtii* is an ideal organism for studying single cell organelle regeneration because it has two linear flagella that grow back upon being cut or shed (1). The kinetics of flagellar growth have been well documented, and much is known about its inner components and growing process, but how the flagellum consistently reaches the same steady-state length is a mystery. Multiple different theoretical models have been developed to explain this robust regeneration, and recent work demonstrated the feasibility of a model in which the length of the flagella is governed by a diffusive gradient across its length (2, 3).

In this study, we further develop the diffusion model by analytically deriving the growth curve as a function of time and the relevant physical parameters. This shines light onto which factors are limiting at different stages in the growth. It also lets us predict steady-state length from observed physical parameters and predict physical parameters from observed steady-state length.

In order to understand the length control model, one must first understand how a *Chlamydomonas* cell builds its flagella. The flagellum is made of nine doublet microtubules, and to get longer, new tubulin (the building blocks of microtubules) must be delivered to the flagellar tip. The mechanism for transporting tubulin to the tip is called intraflagellar transport, or IFT (4–8). In IFT, tubulin and other building materials such as axonemal dynein arms are bound to protein complexes of ∼20 polypeptides called IFT particles. These IFT protein complexes form linear arrays called “trains”, and are pulled to the distal tip by heterotrimeric kinesin-2 motors (9–12). Upon arrival at the tip, the tubulin and other building blocks are added to the flagellum, increasing its length. To counter this length increase, tubulin is continually removed from the flagellar tip at a constant, length-independent rate (13, 14). IFT happens continuously throughout the lifespan of a *Chlamydomonas* cell, and when the rate of IFT-driven assembly equals the rate of length-independent tubulin removal, steady-state length is achieved.

IFT begins through a process called “injection”, in which IFT trains are released from docking sites at the flagellar basal body and transition zone and transported into the flagellum itself (15). Injection is not fully understood, but it appears that IFT material injects into the flagellum from the basal body upon accumulation of motors in the basal body. Quantitative live cell imaging has shown that the rate of injection is a decreasing function of the length of the flagellum (16, 17). This implies some sort of sensing mechanism that allows the basal body to sense the flagellar length. The sensing mechanism here is unknown, and is the core puzzle that length control models try to solve (16, 18, 19). A clue has come from a recent study that showed that kinesin motors diffuse within the flagellum and are not actively transported back to the base (20). In the model explored by Hendel et al., the length-dependent rate of IFT is generated by the kinesin motors diffusing back to the basal body from the tip, using the time it takes to diffuse back as a proxy for length measurement. The longer the flagellum, the longer it takes for kinesins to diffuse back to the base, and therefore the longer it takes for enough kinesins to accumulate in the base to power injection. This explains, in principle, how longer flagella inject less building material per second. The model assumes that kinesins are conserved and not drawn from the cell body in significant number. This would eliminate the need for a currently undiscovered signaling pathway, and would allow the already-known components of IFT to generate its own length dependence. In this study, we will take the conclusions from Hendel et al. and further develop the analytical formalism of the diffusion model to show how altering the diffusion coefficient and IFT velocity would affect observables like steady-state length and regeneration time. We identified three factors that limit flagellar growth at different phases of its regeneration, which lead to two possible rate-limiting steps of flagellar growth at steady state. We then used the upgraded model to attempt to explain observed length changes in length-altering mutants by calculating what changes in diffusion coefficient and IFT velocity are necessary. We arrived at the conclusion that changes in diffusion coefficient may be responsible for the length changes in the mutants.

## 2. Model

We treat the flagellum as a linear track for kinesin motors (Figure 1). The position on the track is labeled by *x* with *x* = 0 corresponding to the base and *x* = *L*(*t*) corresponding to the tip of the flagellum, where *L*(*t*) is the length of the flagellum at time *t*. We distinguish four populations of kinesin motors: (i) motors that actively carry cargos from the base to the tip with a constant velocity *v*; (ii) motors that accumulate at the tip after the delivery; (iii) motors that diffuse back to the base from the tip with a diffusion coefficient *D*; (iv) motors that accumulate at the base when diffusion is completed.

**Figure 1:**
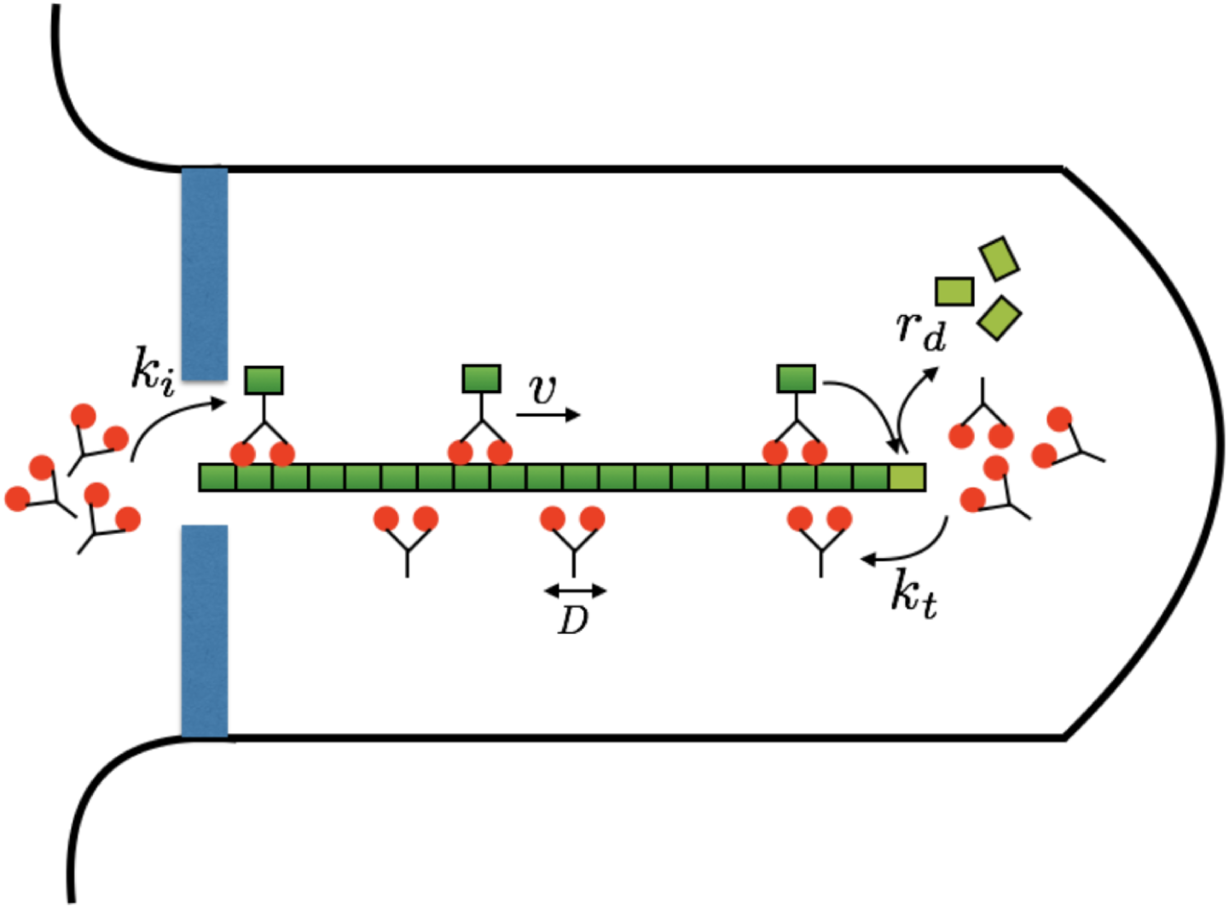
Illustration of the model. Molecular motors carry the building blocks for flagellar assembly from the base to the tip and travel with a constant speed *v*. When reaching the tip, the motors unload the cargo and the flagellum elongates by a unit of *δ*. The motors dwell at the tip and switch to a diffusive state with a transition rate *k*_*t*_. The motors diffuse back to the base with a diffusion coefficient *D* and accumulate at the base, waiting for injection into the flagellum with a transition rate *k*_*i*_. The flagellum has a spontaneous disassembly rate of *r*_*d*_. The total number of molecular motors is assumed to be constant.

The linear number density of active motors *ρ*_*a*_(*x, t*) of type (i) is governed by the equation

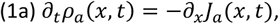

with the convective current

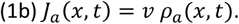

The number of type (ii) motors *N*_*t*_ dwelling at the tip is described by

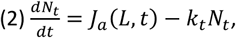

where *k*_*t*_ is transition rate for a motor dwelling at the tip switching to a diffusive state.

The linear number density of diffusive motors *ρ*_*d*_(*x, t*) of type (iii) obeys the simple diffusion law:

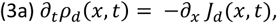

with the diffusive current

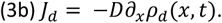

The number of type (iv) motors *N*_*b*_ accumulating at the base is described by

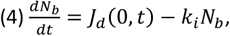

where *k*_*i*_ is the injection rate of motors from the reservoir at the base to the flagellum track. The total number of motors *N* includes all four populations of motors and reads

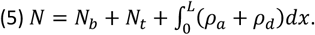

We assume the total number of motors is conserved and this imposes the boundary conditions at the base

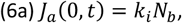

and at the tip

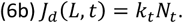

The growth dynamics of the flagellum are governed by the equation

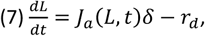

where *δ* denotes the length elongation caused by arrival of a single kinesin motor, and *r*_*d*_ denotes the de-polymerization speed which is independent of the length.

## 3. Results

We can numerically solve the dynamic equations of (1)-(4) and (7) to have the exact growth curve *L*(*t*) for flagellum of length *L* as a function of time *t*. The parameters used in our numerical solutions are listed in Table 1. Due to the small elongation increment *δ*, we can also make a quasi-static assumption that at each length *L*, the spatial distribution of molecular motors reaches the steady state for that particular length *L* (see Appendix A). The analytical results obtained by this quasi-static assumption almost perfectly overlap with the exact numerical solution (Figure 2a, c and e). Therefore, for the rest of the paper we only show results obtained with the quasi-static assumption.

**Table 1:** Parameters of the model.

| Parameters | Description | Reference Value | Varied range | References |
| --- | --- | --- | --- | --- |
| $v$ | Kinesin motor velocity | $2 \mu\text{m/s}$ | $0.1 - 10 \mu\text{m/s}$ | (20) |
| $D$ | Kinesin motor diffusion coefficient | $20 \mu\text{m}^2/\text{s}$ | $0.1 - 80 \mu\text{m}^2/\text{s}$ | (20) |
| $k_i$ | Injection rate of motors at the base | $1 \text{ s}^{-1}$ | $1 \text{ s}^{-1}$ | Arbitrary |
| $k_t$ | Transition rate to diffusive state for motors dwelling at the tip | $0.5 \text{ s}^{-1}$ | $0.5 \text{ s}^{-1}$ | (20) |
| $r_d$ | Spontaneous depolymerization speed of flagellum | $0.004 \mu\text{m/s}$ | $0.004 \mu\text{m/s}$ | (33, 34) |
| $N$ | Total number of motors | 40 | 40 | (13) |
| $\delta$ | Elongation length of the flagellum upon the arrival of a motor at the tip | $0.00125 \mu\text{m}$ | $0.00125 \mu\text{m}$ | (13) |

**Figure 2:**
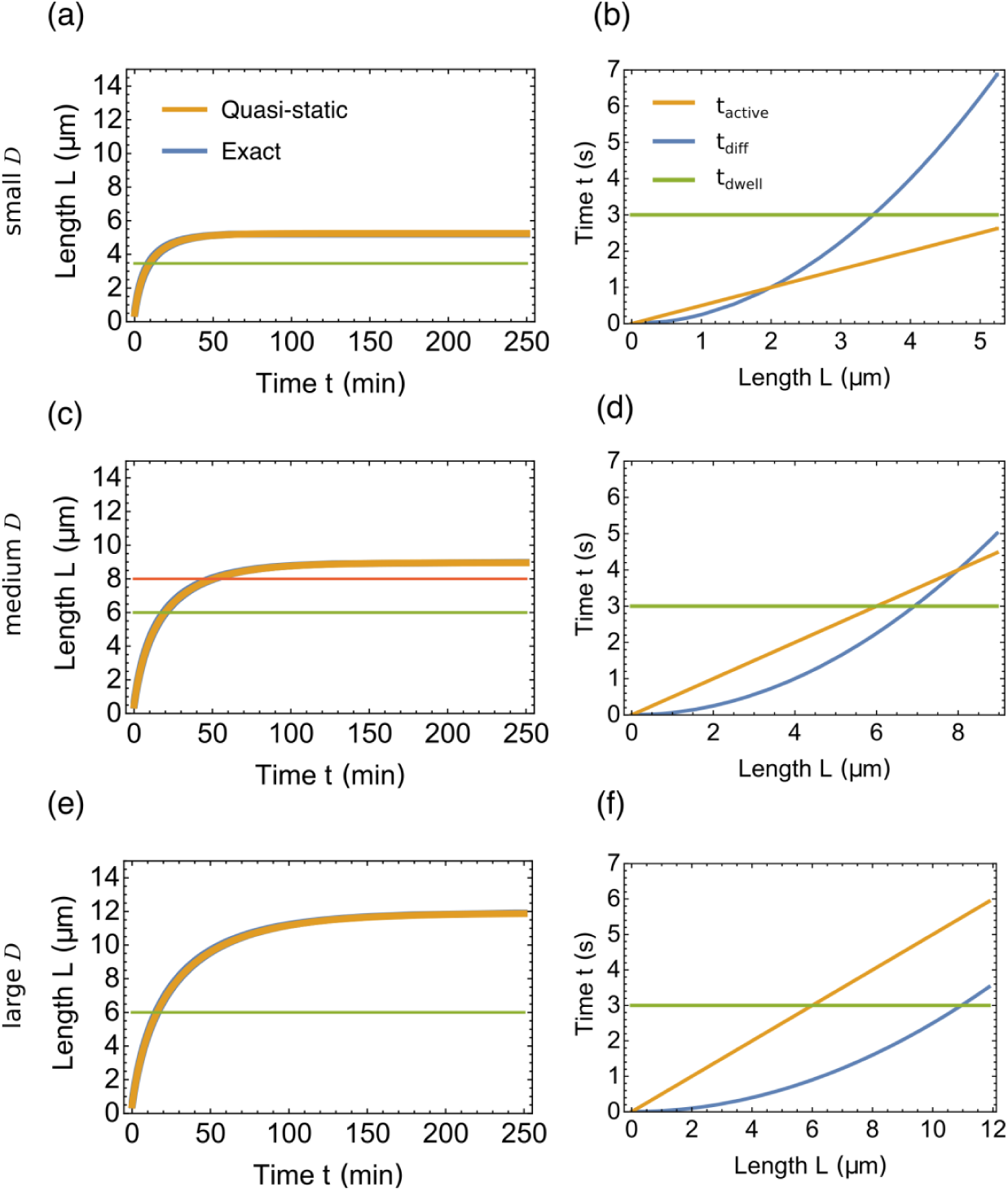
Growth dynamics of the model. (a, c, e) Growth curve of the flagellum for small diffusion coefficient *D* = 2 μm^2^/s in (a), medium *D* = 8 μm^2^/s in (c) and large *D* = 20 μm^2^/s in (e). The blue curve represents the numerical solution, i.e., the exact solution. The orange curve represents the analytical solution obtained by the quasi-static assumption. The two curves almost overlap to the extent that the blue one is invisible. The horizontal lines represent the length at which the rate limiting step changes. (b, d, f) The time a single motor spends on different steps during a transportation-diffusion cycle for the same diffusion coefficient as in (a, c, e). The three curves include *t*_active_ for a motor to travel from the base to tip (orange), *t*_diff_ for a motor to retrieve from the tip to the base via diffusion (blue), and *t*_dwell_ for a motor to dwell at the tip waiting from retrieval and at the base waiting for injection (green).

### 3.1 The rate-limiting step changes as the flagellum grows

Typical growth curves of the flagellum are demonstrated in Figure 2 for three diffusion coefficients. Each growth curve rapidly increases at first and then slowly plateaus to the steady state length. The growth can be divided into different stages based on the rate-limiting step. To see this, we express the growth rate of the flagellum under the quasi-static assumption as

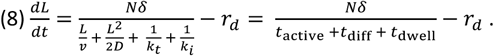

where 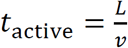 denotes the time for a motor to transport the assembly unit of the flagellum from the base to the tip, 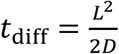 denotes the root-mean-square time for a motor to diffuse back to the base from the tip, and 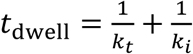 denotes the total time a motor dwells at the base and at the tip. At short length scale, *t*_dwell_ always dominates over the other two time scales, and motors spend most of their time dwelling at the tip and the base (Figure 2b, d and f, green lines). In this regime, the duration that the motor spends traveling between the base of the tip is negligible, so the flagellar growth rate is independent of length. When the flagellum grows longer, either the diffusive time *t*_diff_ dominates if *D* is small (Figure 2b), or the transportation time *t*_active_ dominates if *D* is large (Figure 2f). For an intermediate *D*, the growth is divided into three stages, in which the dominant time scales are *t*_dwell_, *t*_active_ and *t*_diff_ (Figure 2d). Measurements of flagellar growth kinetics have clearly shown that growth rates are constant for flagella shorter than 4-5 micron (1). In a different algal species, Spermatozopsis similis, flagella grow at a constant rate over their whole length, suggesting that in that organism, t_dwell_ is always the dominating factor (20).

### 3.2 Diffusion vs. active transport as the rate-limiting step at the steady state

One would expect that diffusion will always be the rate-limiting step at steady state. This is because the diffusion time *t*_diff_ scales with *L*^2^, while the motor transportation time *t*_active_ scales with *L*. For a sufficiently long flagellum, *t*_diff_ always dominates over *t*_active_. However, the steady state length *L*^*ss*^ might not be long enough to have *t*_diff_ greater than *t*_active_. In Figure 3a, we show the three time scales at steady state as a function of diffusion coefficient *D*. For small *D, t*_diff_ dominates over the other time scales. However, as *D* increases, *t*_active_ becomes greater than *t*_diff_, and the steady state length of the flagellum becomes limited by the active motor transport. If we fix the diffusion coefficient but vary the motor velocity, the growth will change from *t*_active_-dominance to *t*_diff_-dominance (Figure 3B). A phase diagram is shown in Figure 3c. Generally, a larger diffusion coefficient *D* favors motor-limited growth, and a faster motor speed *v* favors diffusion-limited growth.

**Figure 3:**
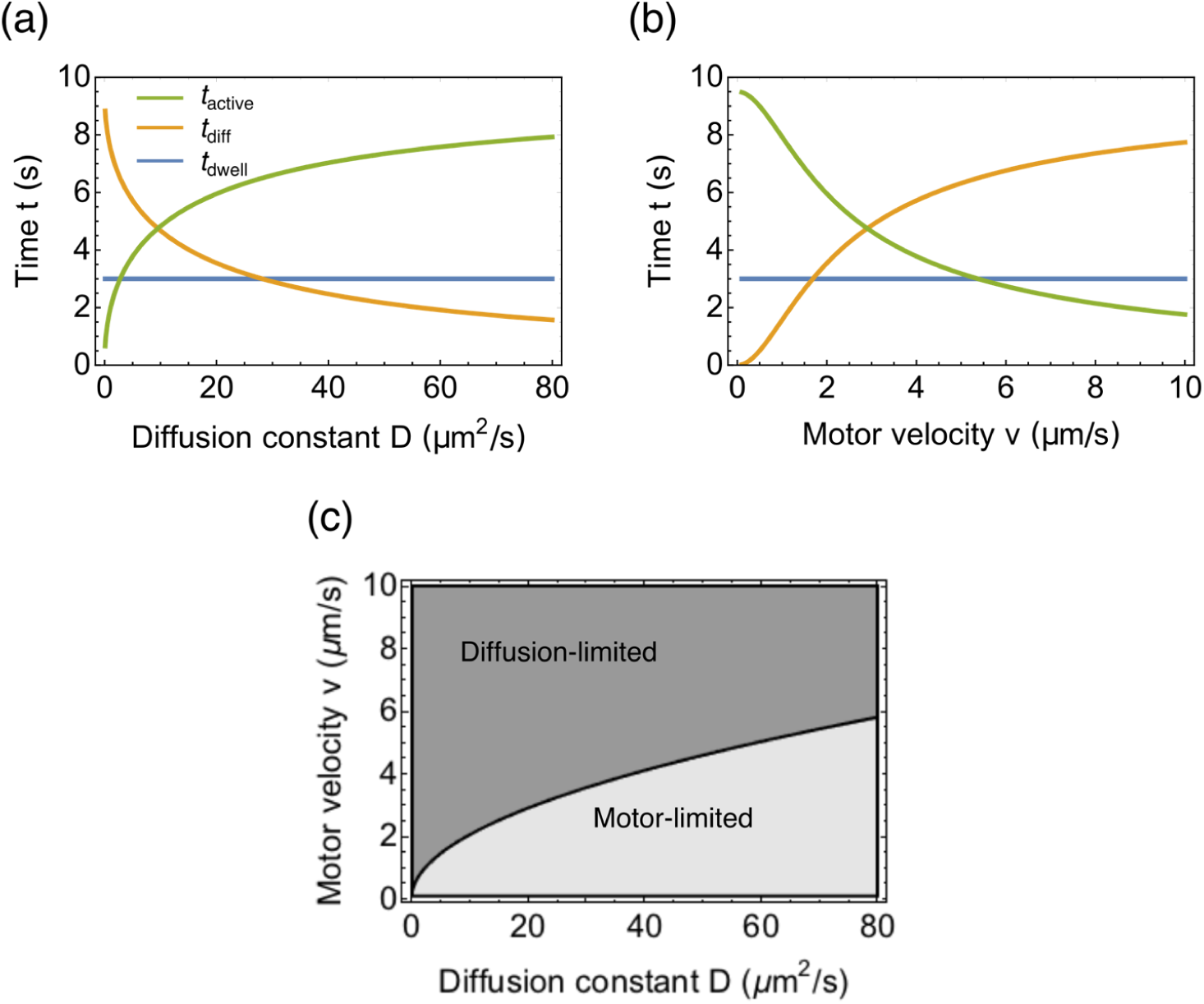
Influence of the motor velocity *v* and diffusion coefficient *D* on the rate-limiting step at steady state. (a) The three time scales as a function of diffusion coefficient *D*. (b) The three time scales as a function of motor velocity *v*. (c) Phase diagram for the rate limiting step at steady state.

### 3.3 A dramatic increase in steady state length *L*^*ss*^ requires a dramatic increase in diffusion coefficient *D* if motor velocity *v* is small

The steady state length of the flagellum *L*^*ss*^ can be obtained by setting 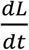 in Equation (8) to zero. This leads to the analytical result

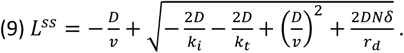

The steady state length *L*^*ss*^ increases with both diffusion coefficient *D* and motor velocity *v*. For a small motor velocity, increasing the diffusion coefficient does not lead to significant increase in *L*^*ss*^ because it is mainly set by the small motor velocity (Figure 4a, green line). For instance, if the motor velocity *v* is 1μm/s and *L*^*ss*^ is 5*μm*, it would be impossible to increase to increase the length to 10 *μm* because even in the limit of an infinitely large diffusion coefficient *D* → ∞, the maximum length *L*^*ss*^ is 9.5*μm*. The analytical proof of this limit is derived in Appendix B.

**Figure 4:**
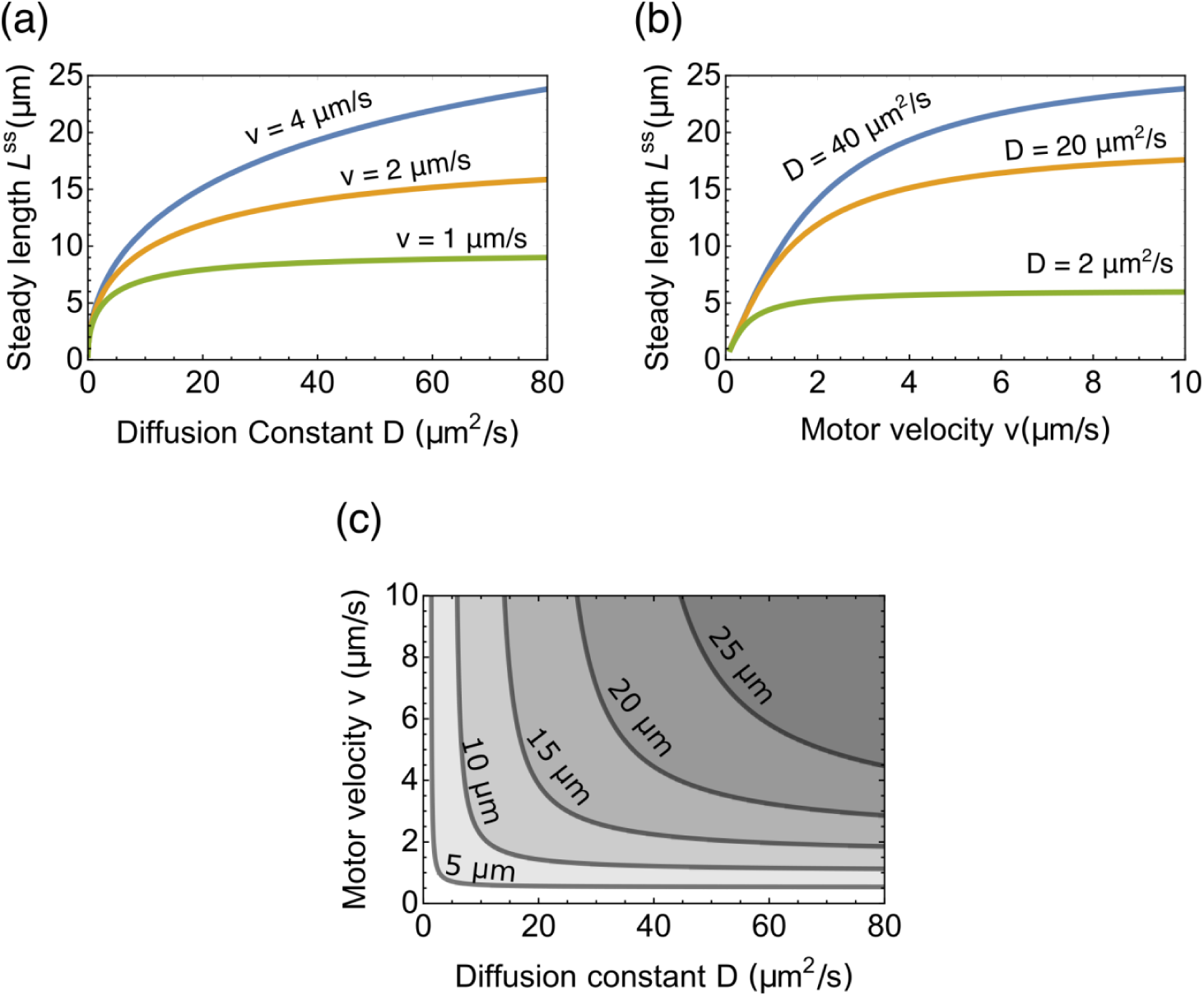
Influence of the motor velocity *v* and diffusion coefficient *D* on the steady state length *L*^ss^ of the flagellum. (a) The steady state length *L*^*ss*^ as a function of diffusion coefficient *D* for different motor velocities. (b) The steady state length *L*^*ss*^ as a function of motor velocity *v* for different diffusion coefficients. (c) The contour plot of *L*^*ss*^ as a function of both diffusion coefficient *D* and motor velocity *v*.

However, if the motor velocity *v* is 2 μm/s, the diffusion coefficient *D* must only increase from 1.8 μm^2^/s to 11.1 μm^2^/s to increase length to ten microns, the typical length of wild type *C. reinhardtii* cells. Similarly, for a small diffusion coefficient, increasing the motor velocity does not lead to significant increase in *L*^*ss*^ either (Figure 4b, green line).

### 3.4 Growing time T of the flagellum increases with motor velocity and diffusion

In this section, we study the time *T* a flagellum needs to grow to its steady state. Mathematically the solution of *L*(*t*) to the mean field equation (8) takes an infinite amount of time to reach steady state length, i.e., *L*(*t*) → *L*^*ss*^ when *t* → ∞. However, as the mean field equation neglects fluctuations of the length at steady state, we define the growing time *T* as the amount of time to reach 95% of the steady state length, i.e., *L*(*T*) = 0.95 *L*^*ss*^. Figure 5 plots numerical solutions of T as a function of motor speed and diffusion coefficient. One might expect that a fast-transporting motor or a fast-diffusive motor will reduce the time to construct a flagellum, but the results show that the growing time *T* increases with the diffusion coefficient *D* and the motor velocity *v* (Figure 5a and b). This is because the steady state length also increases with *D* and *v*. The reduction in time due to increased *v* or *D* cannot compensate for the increased time due to length elongation.

**Figure 5:**
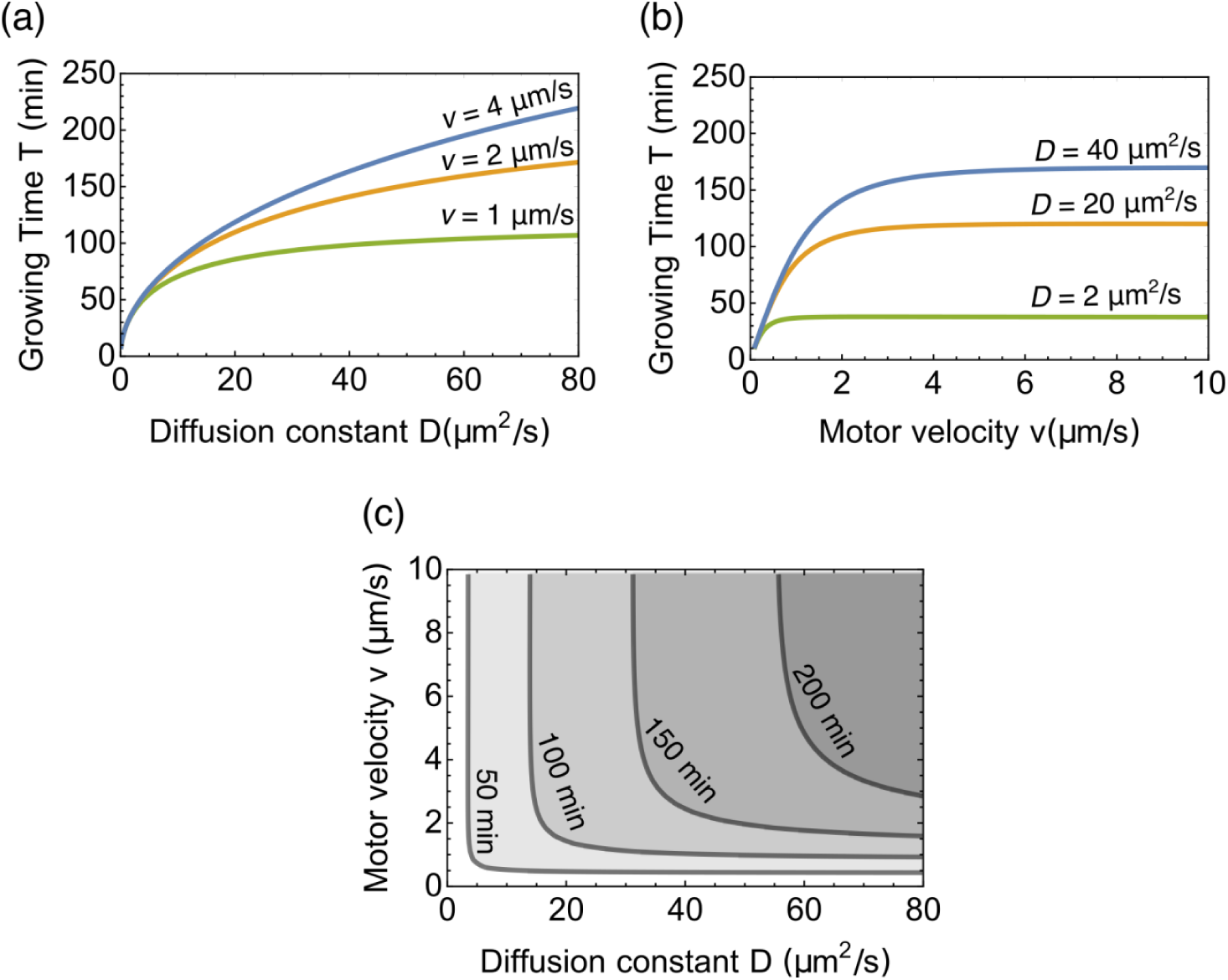
Influence of the motor velocity *v* and diffusion coefficient *D* on the growing time *T* of flagellum. (a) The growing time *T* as a function of diffusion coefficient *D* for different motor velocities. (b) The growing time *T* as a function of motor velocity *v* for different diffusion coefficients. (c) The contour plot of *T* as a function of both diffusion coefficient *D* and motor velocity *v*.

### 3.5 Parameter changes that maintain the steady state length but alter the growing time

One may notice that the contour lines for the steady state length *L*^*ss*^ do not exactly overlap with the contour lines for the growing time *T* (Figure 6a). The implication of this difference is that growth kinetics does not uniquely determine the steady state length, and one can alter the growing time *T* while maintaining the steady state length *L*^*ss*^ constant, or vice versa. A recent experiment found that mutants in ida5 which affect actin, show a slower growth kinetics (i.e. longer *T*) but reach the same steady state length as wild-type. Based on our model, this could be a result of the combination change of reduced motor velocity and enhanced diffusion coefficient (Figure 6b and c). Our model predicts that the change in the growing time is larger for longer flagellar, which can be tested by future experiments.

**Figure 6:**
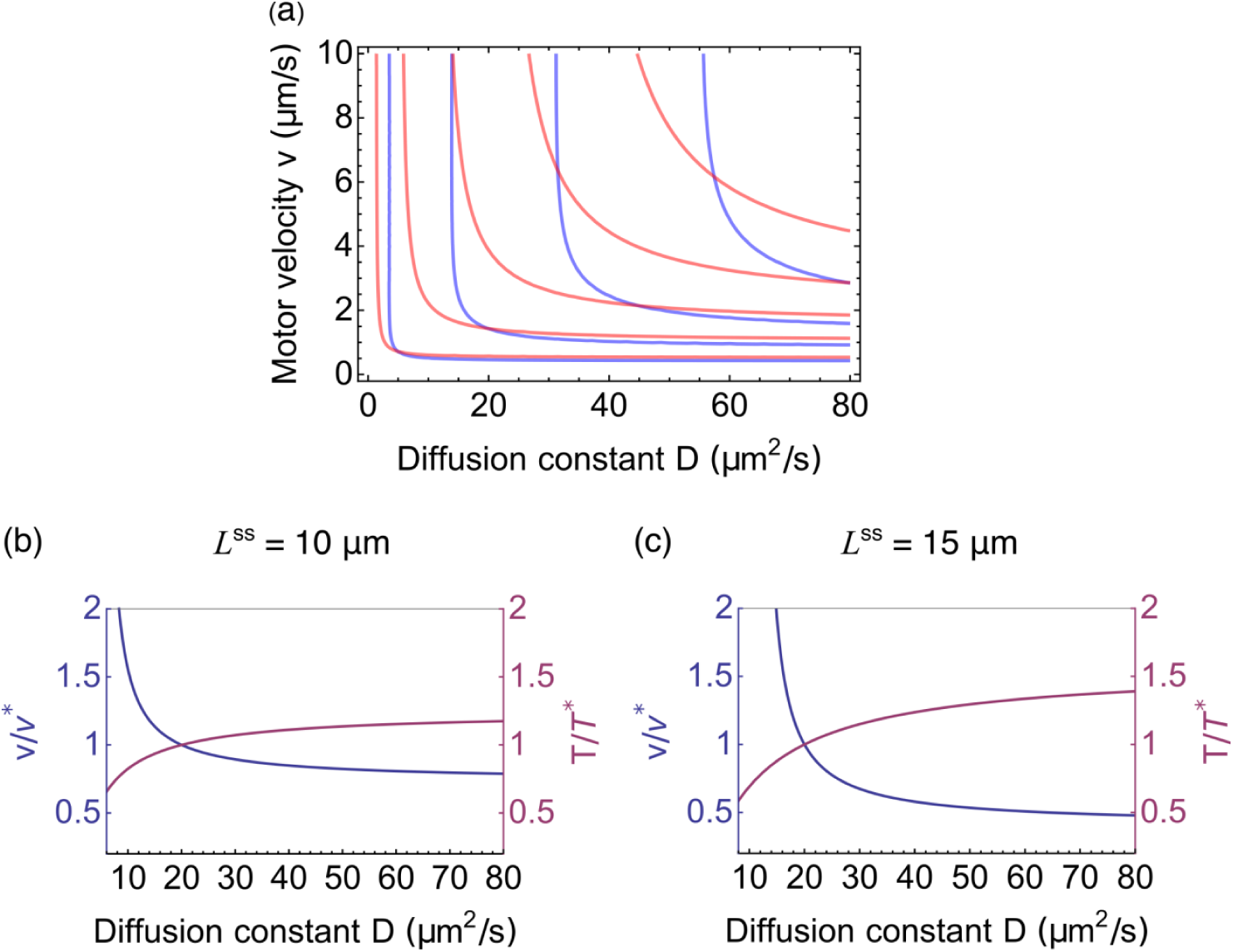
Possible parameter changes that keep the steady state length *L*^*ss*^ constant while altering the growing time *T*. (a) Overlay of the contour plots for growing time *T* (blue) and for steady state length *L*^*ss*^ (red). From left to right, the contours for growing time *T* are 50,100,150,200 min, and for steady state length *L*^ss^ are 5, 10, 15, 20, 25 *μm*. (b, c) Relative change of motor velocity (left axis) and growing time (right axis) as a function of the diffusion coefficient along the contour of *L*^*ss*^ = 10μm in (b) and *L*^*ss*^ = 15μm in (c). The reference velocity is defined as *v*^*^ = *v*(*D* = 20μm^2^/s) and the reference growing time *T*^*^ = *T*(*D* = 20μm^2^/s).

## 4. Discussion

A large part of the explanatory and predictive power of the model is in generating hypotheses to explain length mutants and motivating experiments to test these hypotheses. We can now examine a length mutant, note its length change from wild type, and determine what changes in velocity and diffusion are necessary to achieve the length change. Here we discuss *pf14*, a mutant that is missing the radial spoke head in the flagellum. In wild type *Chlamydomomas*, the two flagella beat in a cyclic pattern resembling a breast stroke: a semi-circular power stroke to swim forward followed by a recovery stroke to return them to their initial position. *pf14*, on the other hand, has paralyzed flagella and cannot swim. What is puzzling about this mutant is that its flagella are about half as long as wild-type (*pf14* mutants are 3-6 μm in length, while wild type flagella are usually of 10-12 *μm* (21). This short flagellar phenotype is common among the group of motility mutants, especially the ones with completely paralyzed flagella (22–27). To our knowledge, no study has explained the connection between paralysis and length decrease – in fact, researchers have viewed intraflagellar transport and flagellar beating as two independent processes. This is not without reason, as beating relies on axonemal dynein and other regulatory and structural components to bend doublet microtubules, components that are not involved in IFT. Even when detached from the cell body, the flagellum equipped with the motility apparatus is capable of producing a high-frequency waveform as long as ATP is provided (28).

While it is possible that the length change comes from a structural instability caused by the mutation, could it instead be because the paralysis of the flagella alters the IFT-diffusion system that could be responsible for length control? All existing measurements of IFT kinetics have been carried out in immotile flagella, either in paralyzed mutants or in wild-type cells whose flagella are adhered to a glass surface. Consequently, there is no experimental information about how IFT kinetics might or might not change in beating flagella compared to immotile flagella. Here we use our model to explore the plausibility of the idea that flagellar beating can influence IFT and thus might act as a wrongfully neglected factor in the length control system.

### 4.1 An increase in diffusion coefficient is necessary for the increase of steady state length

Based on our results, we consider two possible explanations for the significant increase in length in a beating flagellum compared to an immotile one: (i) The motor velocity remains unchanged and the increase is due to enhanced diffusion. (ii) The diffusion coefficient remains unchanged and the increase is due to increased motor velocity. The experimentally measured diffusion coefficient *D* = 1.68 ± 0.04 μm^2^/s and motor velocity *v* = 2.1 ± 0.4 μm/s in a paralyzed mutant (20). With assumption (i), to account for length increase in a beating flagellum from 5μm to 10μm, the diffusion coefficient needs to increase from 1.75μm^2^/s to 10.55 μm^2^/s. With assumption (ii), it is impossible to account for the length increase because even in the limit of infinitely large motor velocity *v* → ∞, the length of the flagellum approaches a maximum of 5.65μm. Therefore, an enhanced diffusion is necessary and sufficient to account for the observed length increase. In any case, there is no plausible way that flagellar beating would alter the velocity of the motor. However, we can imagine several ways that beating could alter the diffusion coefficient of kinesin, which we will consider in turn.

### 4.2 Centrifugation effect of kinesin motors

The first mechanism we considered was inspired by the experimental observation that kinesin-2 is less dense than the flagellar matrix and floats to the top when a matrix preparation is centrifuged at high speed (H. Qin unpublished data). Based on this observation, we consider a model in which the roughly circular beating of the flagellum was enough to cause a significant centripetal force on the kinesin motors back towards the base, speeding up the diffusive return time. To model this scenario, we approximated the flagellum and its beating as a cylindrical rod revolving around one of its ends like the hand of a clock. The contents of the cylinder will experience a centrifugation effect, and the kinesins will move towards the base if they are less dense than the surrounding solution. While this is not equivalent to increasing the diffusion coefficient, it is an increase in the speed of diffusive return. Approximating the beating as a circular motion will exaggerate the centripetal force because the recovery stroke of the beating does not have the same circular appearance as the power stroke. To estimate the magnitude of this effect, we solved the equation for centripetal force to obtain the drift velocity:

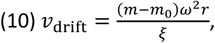

where *m* is the mass of kinesin, *m*_0_ is the mass of the solution displaced by the motor, *ξ* is the friction coefficient (equal to *kT*/*D* where *k* is Boltzmann’s constant, *T* is the temperature, and *D* is the diffusion coefficient), *ω* is the rotation rate, and *r* is the length of the rod. Plugging in the relevant values *D* = 2 μm^2^/s, *kT* = 4.1 pN · nm, *m* = 0 (extreme case where kinesins are massless, to give the maximum possible effect), *m*_0_ = 4 × 10^−22^ kg, *ω* = 300 rad/s, and *r* = 10μm, we get that the drift velocity *v*_*drift*_ is on the order of 10^−7^ μm/s, which means it would take three years for the kinesins to get from the tip to the base with this effect alone. If we translate the time sped up by the centrifugation drift into diffusive time, it amounts to an effective diffusion constant of *D*_eff_ which satisfies

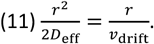

The effective diffusion constant increase *D*_eff_ is only on the order of 10^−6^ μm^2^/s, which is negligible compared with measured value of *D*∼2μm^2^/s. We can therefore rule out the centrifugation effect as a means of generating any substantial length increase upon beating.

### 4.3 The increased diffusion coefficient in a beating flagellum might be explained by shear-thinning

An alternative way that flagellar beating could influence diffusive return of kinesin is via shear of the flagellar matrix (Figure 7a). If we think of the flagellum as an elastic rod, when it is bent, parts of the rod are stretched and parts are compressed. The maximum shear displacement Δ can be calculated as (29).

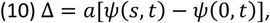

where *a* is the radius of the rod, and *ψ*(*s, t*) is the tangent angle along the arclength *s* at time *t*. The corresponding shear rate is

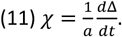

We select 7 frames in a periodic beating cycle of a flagellum and calculate the shear displacement and shear rate by measuring the tangent angle at equally spaced points along the arclength of the flagellum (Figure 7 b-f). In a beating period of *T* = 0.014 *s* (30), the variation of the shear displacement *δ*Δ ≡ max(*Δ*) − min(Δ) is typically around 0.4 *μm*. Here the max and min are taken with respect to the time *t* in a period. The periodic variation of the shear displacement can induce a shear flow in the matrix and move the motor back and forth, therefore creating an effective diffusion. Assuming the motor completely follows the induced shear flow, the distance travelled by the motor in a period is *δ*Δ. The effective constant 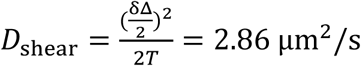, which is not enough to account for the required increase in diffusion coefficient from 1.75μm^2^/s to 10.55 μm^2^/s. Furthermore, this analysis tends to overestimate the diffusion coefficient as it assumes the maximum shear Δ at the boundary is completely transferred to the diffusive motion of motors, which apparently neglects the gradient of the velocity field and dissipation of the energy transfer. We conclude that shear is not large enough to increase diffusion constant significantly by means of an advective mechanism. Could shear have any other effect?

**Figure 7:**
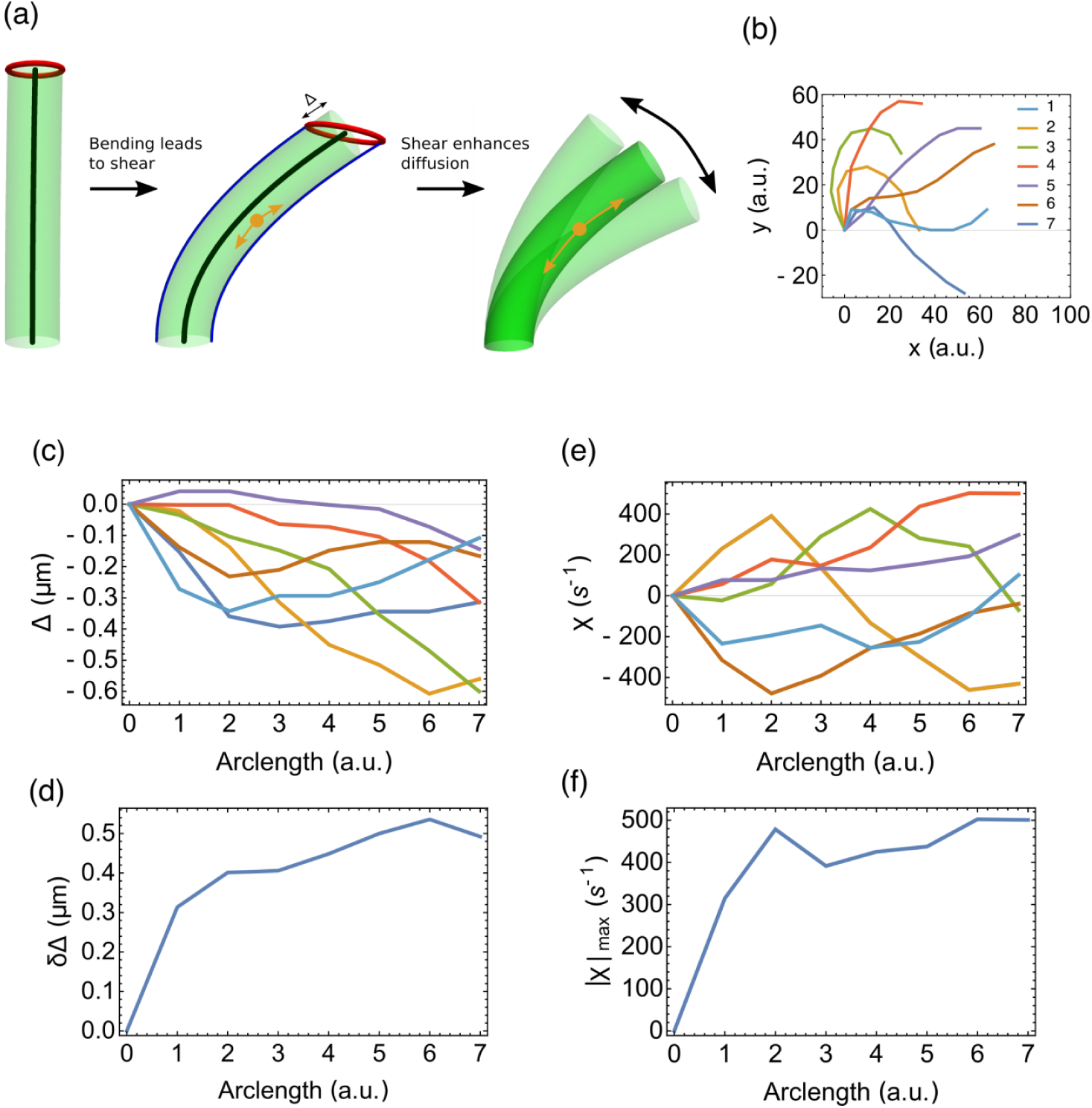
Beating of flagellar leads to enhanced diffusion of motors. (a) The flagellum is depicted as a rod. Bending of the rod leads to stretch on one side, and compression on the other side. The two blue curves represent curves on the rod’s surface that have the same length as the central axis (black line). The red circle represents all the end points on the rod’s surface that have the same length with the central axis. The shear induced by periodic beating of flagellum can enhance the diffusion of molecular motors via the shear-thinning mechanism, thus increasing the length of flagellum compared with paralyzed mutants. (b) Selected beating shapes of a flagellum in a beating cycle. The number indicates the order of the sequence. (c, d) Shear displacements Δ in (c) and its variation *δ*Δ in a beating period in (d). (e, f) Shear rates *χ* in (c) and its maximum in a beating period in (f).

It is well known that solutions made of soft polymers become less viscous under shear deformation. This effect is known as shear thinning. In an equilibrium solution, the diffusion coefficient *D* of a particle and its friction coefficient *ξ* obey the Einstein relation *ξD* = *k*_*B*_*T*, where *k*_*B*_ is the Boltzmann constant, and *T* is the absolute temperature. Because the friction coefficient *ξ* is proportional to the viscosity *η*, we would expect the product of the viscosity *η* and the diffusion coefficient *D* is also a constant. Therefore, a reduction in viscosity *η* caused by shear-thinning might account for the increase in diffusion coefficient *D*. Based on our measurements, the maximum shear rate |*χ*|_*max*_ ≡ max(|*χ*|) of the flagellum is around 600 s^−1^ (Figure 7F). The onset shear thinning rate for biopolymer solutions depends on many factors, such as protein concentration, temperature, ionic strength and even the geometry of the container. The typical onset shear thinning rate for a polysaccharide solution is ∼10 *s*^−1^ and the reduction in the viscosity can be orders of magnitude (31). Recent work on bioink (alginate plus cellulose) shows that the shear thinning effect takes place at very low shear rate (32). Therefore the shear magnitude is large enough to potentially cause shear thinning in the matrix of flagella, and this effect may contribute to enhanced diffusion by reducing the viscosity. Our results thus suggest a novel hypothesis to explain the link between flagellar motility and length, namely, that paralyzed mutants have shorter length because the diffusion constant for kinesin is decreased due to a loss of shear thinning in the flagellar matrix. Our modeling results suggest a need for future experiments to measure viscosity inside the matrix.

## Appendix

### A. Derivation of the growth rate (8) under quasi-static assumptions

In physiological conditions, the length elongation of flagellum is much slower than the motor transportation-diffusion cycle. This is reflected in the small elongation unit *δ* in Eq.(7). We can therefore make the quasi-static assumption that at any fixed length *L*, the distributions of the four populations of motors reach steady state for that particular *L*. This implies that all the time derivatives in Eqs. (1-4) become zero. The distribution of the active motors (i) is homogenous over the flagellum track, and the constant density reads

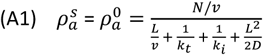

For the diffusive motor, the spatial distribution shows a gradient and reads

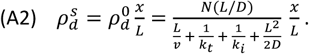

For the motors accumulated at the base, the number

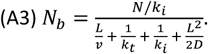

For the motors accumulated at the tip, the number

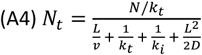

Substituting Equation (A1) into (7), we obtain (8), which is the key equation of our discussion for the dynamics of flagellum growth.

### B. Derivation of the steady state length in the limit of large diffusion coefficient

Denoting 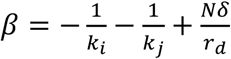, we can rewrite Eq. (9) as

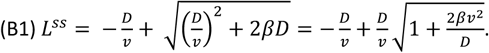

In the limit of *D* → ∞, we can invoke the Taylor series (1 + *x*)^*k*^ = 1 + *kx* + *O*(*x*^2^) for |*x*| ≪ 1.

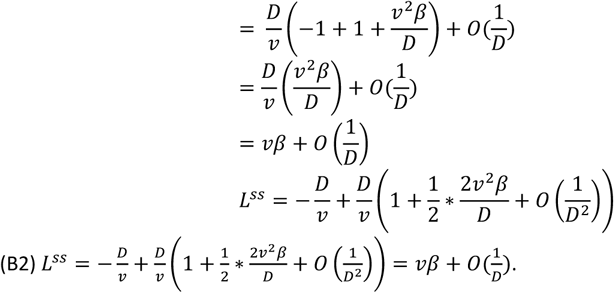

Therefore *L*^*ss*^ → *v²* in the limit of *D* → ∞.

## Author Contributions

RM, NLH, WFM, and HQ designed research. RM and NLH performed research and contributed analytic tools. RM, NLH, WFM, and HQ analyzed data. NLH, RM, HQ and WFM wrote the manuscript.

## Acknowledgments

This work was initiated, and initial stages of the model developed, at two Cell Modeling Hackathon events supported by NSF grant MCB-1411898. NLH and WFM acknowledge support of NIH grant R35 GM130327.

